# Lowering the switching cost related to the activation of burdensome gene circuits promotes cell population homogeneity and productivity

**DOI:** 10.1101/2024.10.14.618176

**Authors:** Lucas Henrion, Vincent Vandenbroucke, Juan Andres Martinez, Julian Kopp, Samuel Telek, Andrew Zicler, Frank Delvigne

## Abstract

The activation of gene circuits can impose a significant burden on cells, leading to heterogeneous expression and reduced productivity. In this work, we focused on the T7 production system in *E. coli* BL21, a prime example of a burdensome gene circuit, to investigate the main cause for this gene expression heterogeneity and methods to mitigate it. Based on continuous cultivation analyzed and control by automated flow cytometry, we quantified the trade-off between cellular growth and gene expression and tracked the cell-to-cell heterogeneity in gene expression (measured as entropy). We concluded that the growth reduction associated to the activation of the burdensome gene circuit, i.e., the switching cost, is at the origin of the population heterogeneity. The loss of growth rate imposed by the burdensome activation of the gene is compensated at the population level by the overgrowth of less induced cells that safeguard the population by generating entropy. We tried to homogenize the population by pulsing the inducer with increasing frequency but found that the population escapes control through promoter mutation, leading to a genotype exhibiting reduced gene expression, but also, reduced entropy. To engineer a more homogeneous population without sacrificing gene expression, we decreased the switching cost associated to the induction by lowering the quality of the main carbon source. This strategy successfully led to a more homogeneous and productive population. Our approach allows for a precise quantification of the trade-off between growth and gene expression in cell population cultivated under dynamic conditions and highlights the importance of the switching cost for designing efficient approaches of cell population control.

## Introduction

Even in a uniform environment, genetically identical cells still display significant differences in gene expression^**2,19,20**^.This heterogeneity stems from two primary sources, i.e., intrinsic noise, which is an inherent characteristic of biochemical reactions, and extrinsic noise, which arises from variations in the content of macromolecules between cells^**1**^. This noisy expression can have functional consequences at the population level by generating phenotypically distinct sub-populations further improving population fitness by providing robustness against sudden environmental changes^**3,22,24**^.

The beneficial impact of phenotypic heterogeneity on population fitness has been reported several times for microbial populations evolving in a natural context^**9,10,11**^. However, in bio-production, cell-to-cell heterogeneity is still perceived negatively^**12,14**^. Indeed, within cell populations engineered for bio-production of recombinant molecules, important cell to cell variability in production and viability was observed^**13,15,26,18**^. To better control the outcome of gene expression, synthetic biology approaches have been used to engineer more robust and predictable gene networks^**4,5**^, yielding more homogeneous induction. Despite all these efforts, the impact of cell diversification dynamics on bio-process performances is still an open question. In this context, our previous work suggests that the characteristics of the gene circuit alone are insufficient to explain the observed degree of heterogeneity in gene expression^**6**^. Instead, analysis of the heterogeneity in gene expression across diverse gene circuits in multiple model organisms revealed that heterogeneity increases with the switching cost associated to the activation of the gene^**6**^. In that work, which focused on the study of cellular heterogeneity in continuous cultivations, switching cost was defined as the reduction in growth rate associated with the activation of a gene circuit. This definition is analogous to production load or metabolic burden and highlights the importance of this previous observation in the context of bio-processes.

To control gene expression, it was proposed that the inducer could be pulsed instead of being added continuously in the cultivation device^**7,21,23**^. The efficiency of this approach was previously confirmed with a optogenetics where the expression of a fluorescent protein was more homogeneous across the population when applying light as pulses instead of continuous supplied^**7**^. It was suggested that pulsatile induction homogenizes expression because the transcription factors expression itself is bursty^**8**^. Previously, we also observed the benefit of pulsing the inducer instead of feeding it continuously, but only for gene circuits exhibiting a important burden^**6**^. This current work aims to shed light on why the expression of burdensome genes is more heterogeneous than non-burdensome genes and to propose new approaches for promoting the homogeneous activation of such genes. To this end, the activation of the T7 system in *E. coli* BL21 (DE3) was selected as a case study for our analysis since its activation results in a significant burden^**6**^ and is characterized by an unstable induction in continuous cultivations^**16**^.

## Results

### The increase in cell population entropy reflects a trade-off between growth and gene expression at the single cell level

To understand the origin of the heterogeneous expression of burdensome genes during continuous cultivation, the T7 expression system in *E. coli* BL21 containing the pET28:GFP plasmid was periodically stimulated to characterize the induction and relaxation phases. The activation of the T7 system was followed by the expression of EGFP, the production both mimicking the expression of a heterologous protein in a bio-process while being easily quantified at a single cell resolution with a flow cytometer (FC). For this purpose, the strain was grown in a Segregostat, where glucose is supplied continuously while lactose (the inducer) is conditionally added in pulses (**Figure 1a**). These pulses are triggered when FC analysis indicates that less than 50% of the population exceeds a pre-set fluorescence threshold that is much greater than the basal auto-fluorescence of the strain (here 1,000 f.u.). The fluorescence data, collected in the FL1-A channel of the FC, can be combined to create a visual representation of expression levels over time. This is achieved by concatenating the data from multiple analysis cycles, resulting in a time density plot that provides a comprehensive overview of the expression patterns (**Figure 1b**). The fluorescence values in the time density plot can be binned, each bin containing a proportion of the population. This binning allows for the quantification of heterogeneity using a metric derived from information theory, Shannon entropy. Unlike the standard deviation or Fano factor, the entropy is not biased by the mean and does not assume a normal distribution. Measured in bits, entropy increases with the degree of cell-to-cell heterogeneity in gene expression. The second type of data that can be extracted from the time density plot is the growth rate of cells as a function of gene expression intensity. This can be achieved by analyzing the fluorescence decrease of each quartile during a relaxation phase (**SI Note 1**). This decrease in florescence is driven by growth and thus this approach quantifies the trade-off between gene expression (fluorescence level) and growth (**Figure 1c**). For the T7 system, gene activation can lead to a dramatic loss of growth rate, termed switching cost, of more than 80% of the initial growth rate.

**Figure 1.**
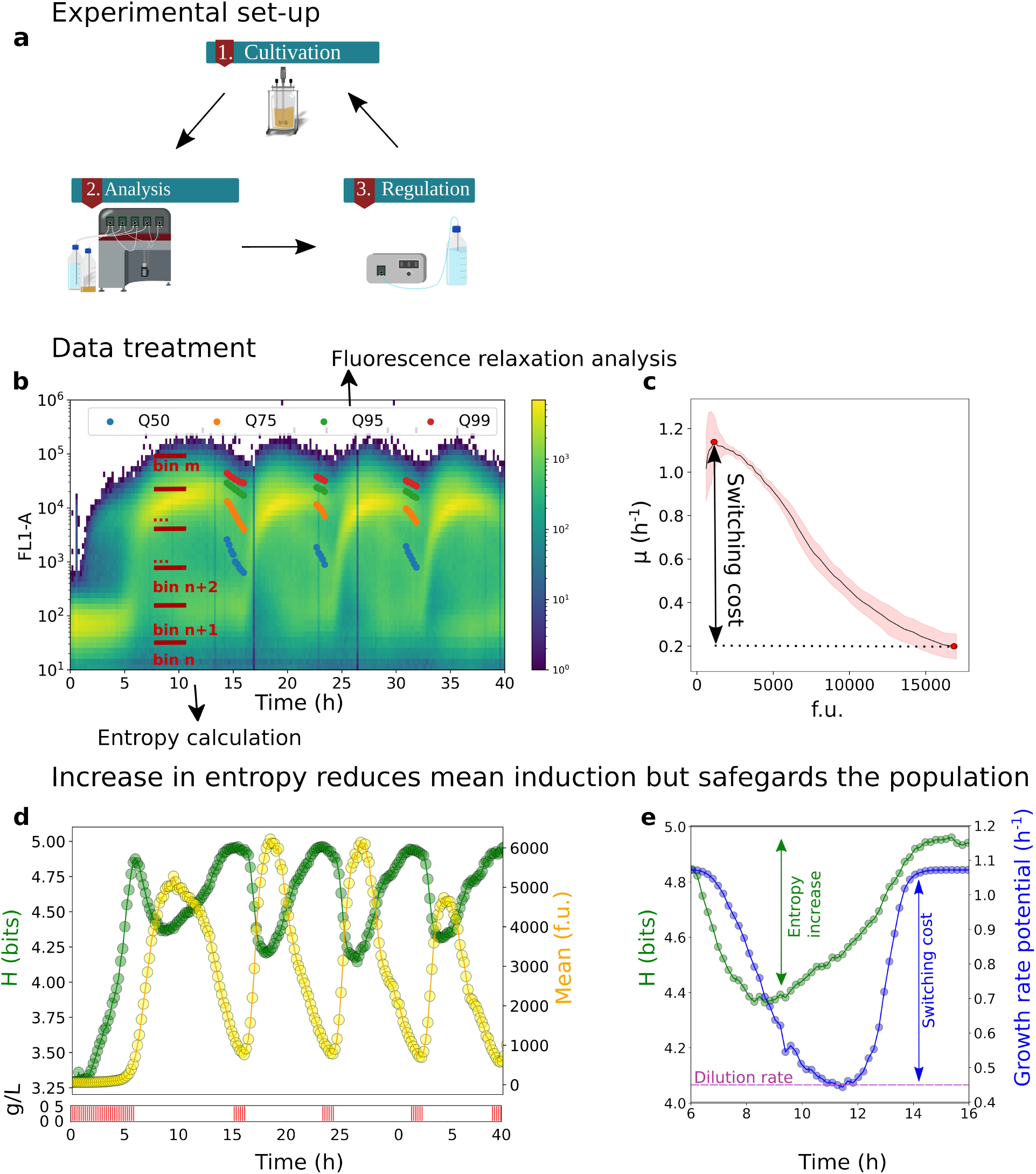
Heterogeneity in the expression of a burdensome gene circuit derives from a burden entropy compensation mechanism: **a** The Segregostat is a cell-machine interface allowing to draw automatically samples from a cultivation device (in our case, a continuous bioreactor) and trigger FC analyses. The collected data are then compared to a pre-set fluorescence threshold that triggers the regulation. **b** The collected data (20,000 cells per analysis) is plotted as a time density plot with the fluorescence on the Y axis (here FL1-A to measure GFP abundance) and the time in hours on the x-axis (two biological replicates **SI Figure 1**, n=2). The y-channel values are binned to compute entropy (H) as a quantitative proxy for heterogeneity. The fluorescence relaxation can be analyzed by measuring the decrease in quartile values. **c** Looking at three relaxation phases and assuming the GFP gets solely diluted by growth, the growth rate associated to the GFP content can be computed (slope of the decay the log10 values of fluorescence quartiles). This reveals that a higher initial fluorescence corresponds to a much lower growth rate. **d** The mean fluorescence increases after each pulsing (0.5 g of lactose per pulse in a 1 l reactor with a OD of 5 as shown by red v-lines below the main graph). H decreases when pulsing and goes up during relaxation. **e** The activation of the burdensome gene reduces the maximum growth rate achievable by the population because of the switching cost. The maximum growth rate achievable by the population was computed from the median fluorescent content and corresponding growth rate gathered from the switching cost curve (c) it is associated with. The population growth rate is nevertheless limited by the dilution rate. Before the population growth rate drops below the dilution rate and the population gets washed-out, the cells with a lower induction overgrow the other highly induced ones and compensate for their low growth rate. We can observe here that growth is restored upon population diversification (measured based on entropy).

With this quantitative metric of heterogeneity, entropy, we analyzed the dynamics of cell population in the Segregostat. As expected, the mean fluorescence rises during induction (following lactose addition) and falls during the relaxation phase, until the next pulse (**Figure 1d**). More intriguingly, the observation of entropy reveals that the population becomes more homogeneous during induction and spreads out during relaxation (**Figure 1d**). Further investigation demonstrated that the increased heterogeneity during relaxation was driven by the trade-off between growth and gene expression (**Figure 1c**). Specifically, cells that are less induced grow faster than those that are more induced, their GFP content decreases faster, and the population gets more heterogeneous. To sum up; the activation of a burdensome gene threatens the population survival in the reactor since if all cells were to have the same degree of activation, the population would be washed out. But the cells with a lower expression of the gene compensate for this burden by outgrowing the others and rescuing growth. This phenomenon is termed Burden-Entropy (B-E) compensation (**Figure 1e**).

### Reducing cell population entropy based on forced induction cycles gives rise to mutational escape

These initial data show that the trade-off between growth and gene expression promotes the heterogeneity of the population. But there also point out that applying a sharp induction phase by pulsing lactose can transiently reduce entropy. We therefore hypothesized that by progressively increasing the induction frequency, the population could be uniformly driven towards higher fluorescence (f.u.) values and reduced entropy (H). Ultimately, this strategy should result in a washout when the population becomes sufficiently homogeneous, such that the median fluorescence 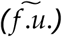 leads to a growth rate of the population lower than the dilution rate (D) imposed by the cultivation device. According to this hypothesis, we considered three cell population outcomes depending on the induction frequency (**Figure 2a**).

**Figure 2.**
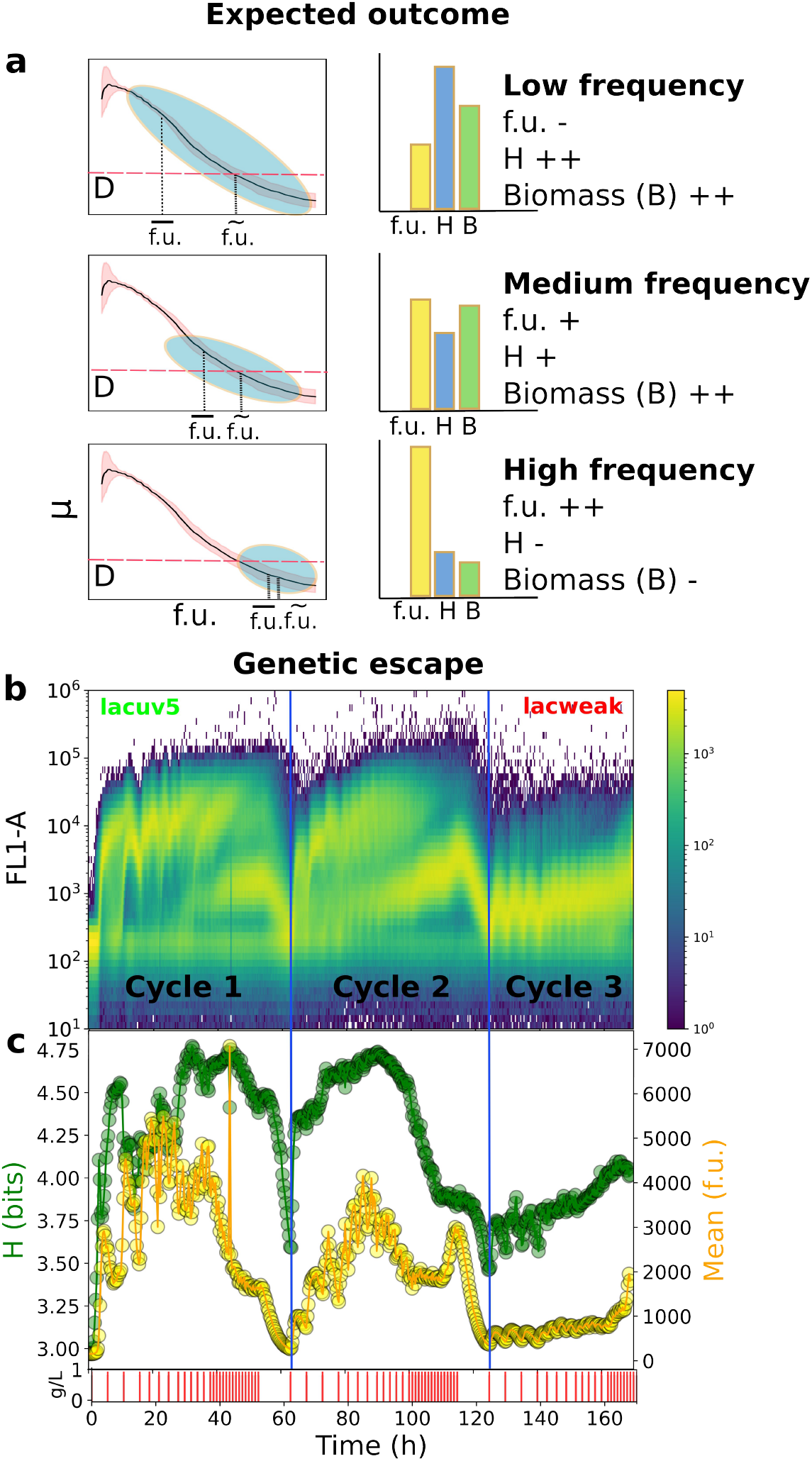
High pulsing frequencies fail at lowering entropy and increasing induction level: **a** The population growth rate corresponds to the growth rate of a cell with a median fluorescence content 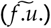. Following the B-E compensation theory, at low pulsing frequency, a wide diversity of induction (blue shade on the graph) is present with a median induction 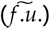 above the mean 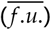. When increasing the pulsing frequency, the median should not increase but the mean induction should go up, resulting in a lower heterogeneity (entropy). It was speculated that as the pulsing frequency increases further, ultimately the population would be washed out because the burden would be too high 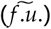 so high that the population growth rate would fall bellow the dilution rate). **b** Unexpectedly, the time density plot (20,000 cells per analysis) shows that an intermediate population appears and that the population does not washout throughout the three cycles of increasingly high pulsing frequency (two biological replicates **SI Figure 2**, n=2). **c** While at first the median induction increased (during the first cycle), it quickly starts going down whilst the entropy increases. Once the intermediate population becomes more prevalent, the entropy is much lower but the mean induction as well.

We hypothesized that at low pulsing frequencies, the B-E compensation is the main mechanism influencing the population dynamics, rescuing the growth rate loss associated with the harsh induction. As a result, the growth rate of the population is determined by the dilution rate imposed on the population and the population remains highly heterogeneous. Then, as the periodic homogenization of the population increases with higher pulsing frequencies, B-E has less to enable population diversification, and thus entropy decreases to the benefit of the mean induction. At a critical point, if the share of the population with high induction is too high. There is not enough heterogeneity to compensate the burden imposed on the population, and the system is washed out.

We then experimentally tested our hypotheses in a continuous culture where increasing frequencies of lactose pulses were applied (**Figure 2b**). We applied three successive cycles of increasing pulsing frequency to the cell population. Unlike the predictions (**Figure 2a**), the cell population never washed-out. During the first pulsing cycle, the mean fluorescence increased upon increasing the lactose pulsing frequency, but the entropy remained quite constant over time (**Figure 2c**). The entropy dropped only for the highest lactose pulsing frequency, but mainly due to the appearance of a new sub-population exhibiting a lower gene expression. The appearance of this sub-population was not expected, and thus before applying a second cycle, the population was left unstimulated during five retention time to clear out any potential memory effect. Despite this time, as soon the pulsing sequence resumed, the intermediate sub-population appeared again and increased in abundance over time before taking over during the third cycle (**Figure 2b**). This sub-population appeared to have lost in induction strength but was more homogeneous. Sequencing revealed that the strong lacuv5 promoter that drives the expression of the polymerase T7 had mutated back to the weaker lac wild-type promoter. This mutation had previously been described and is known to lower the expression of the polymerase, leading to a reduction of the switching cost associated with gene expression^**17**^. Again, this data suggests that the trade-off between growth rate and gene expression greatly impacts the entropy exhibited by the whole cell population.

### Reducing the switching cost promotes population stability and productivity

The initial effort to promote higher population induction by reducing heterogeneity led to mutational escape, an irreversible outcome that has been reported several times as a major cause of decrease in productivity for various bio-processes^**18**^. However, this outcome showed another path to promote a more homogeneous population, reducing the growth rate loss associated with the activation of the gene circuit, i.e, the switching cost. In essence, the mutation reduced the burden as it resulted in a weaker induction. Thus, the mechanism of burden entropy compensation is weakened because there the gap growth rate gap between the induced an un-induced cells is lower.

To promote a more homogeneous population without losing growth potential, we thus need to reduce this growth rate gap between the induced cells and the others without reducing the promoter strength. Reducing the dilution rate imposed on the system should not promote a more homogeneous population because it does not impact the switching cost and thus the competition within the reactor would remain the same. Thus, we propose to reduce the maximum achievable growth rate of the cells. Since the switching cost is the difference between the maximum growth rate of the non-induced cells and the growth rate of the induced one, this option is the only way to reduce the switching cost without reducing gene expression strength. We screened the maximum growth rates of *E. coli* on various alternative carbon sources (**SI Figure 4**). Among the eight carbon sources tested, arabinose and xylose were selected. Arabinose reduced the maximum growth rate by 15% and xylose by 50% compared to glucose, while both maintaining growth rates above the dilution rate used in continuous cultivations of this work.

We then assessed the impact of this strategy on population induction by conducting chemostat experiments, continuously co-feeding lactose with glucose (1 g of lactose per 5 g of the other carbon source), arabinose and, finally, xylose (**Figure 3a**). As hypothesized, cell population induction exhibited lower entropy when xylose was the main carbon source by comparison with glucose (**Figure 3b**). Additionally, unlike the previous pulse-based approach to homogenize the population, this strategy reduces the trade-off with production, resulting in a significantly higher average induction with xylose than with glucose, as reported based on the mean fluorescence level (**Figure 3c**).

**Figure 3.**
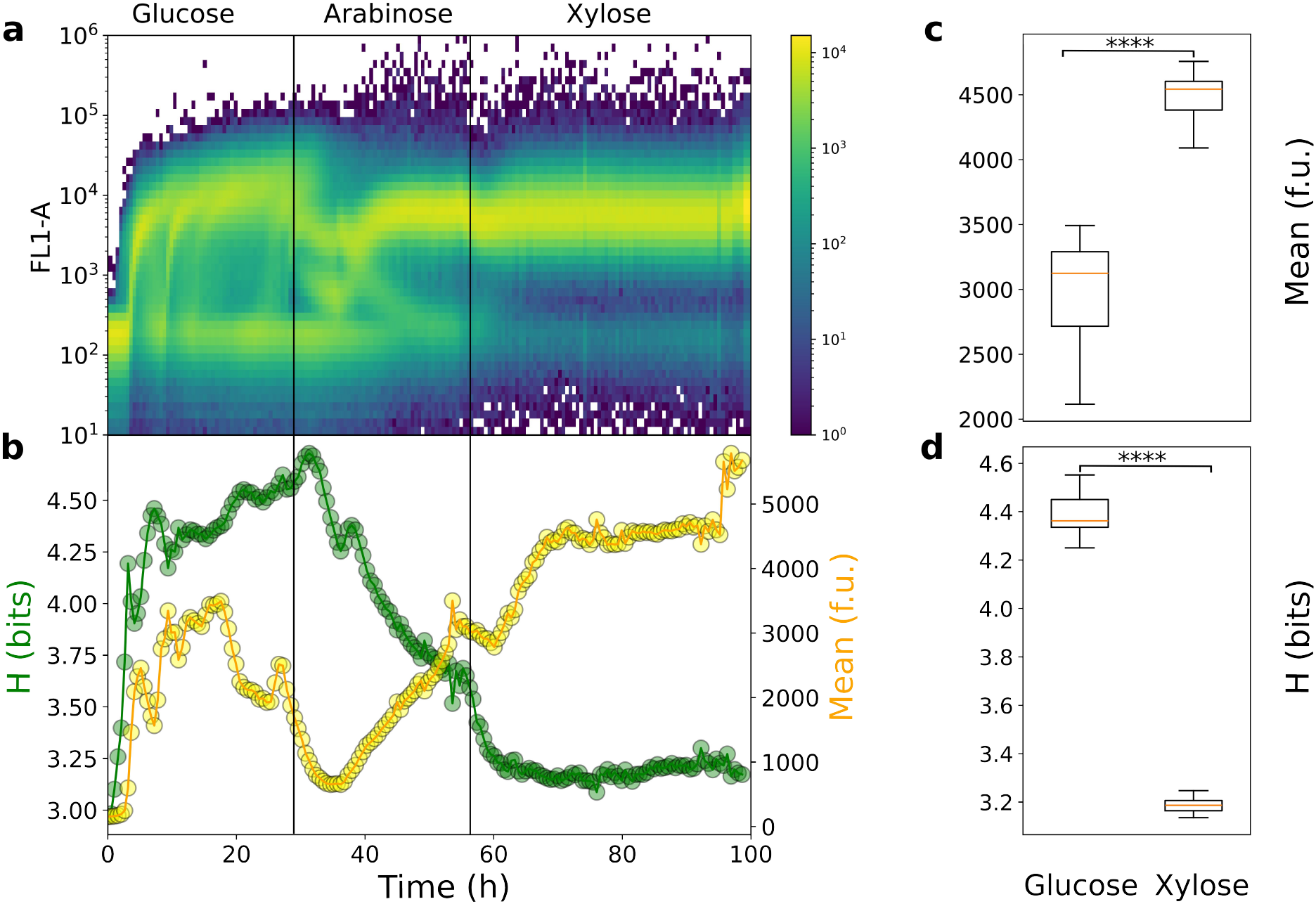
Reducing the global maximum growth rate of the cell population lowers entropy and increases induction: **a** In chemostat cultures, the times-scatter plot (20,000 cells per analysis) shows that the population is highly heterogeneous when grown with glucose as the main carbon source. However, when switching to arabinose, and then to xylose, the population is more homogeneously induced (three biological replicates **SI Figure 3**, n=3). **b** The mean induction increases simultaneously with the decrease in entropy, as shown in further analysis. **c** Cell-to-cell heterogeneity in gene expression (measured as information entropy) is significantly (p<0.001) smaller when xylose is used as the main carbon source compared to glucose. **d** Decreased entropy leads to a significantly (p<0.001) greater mean induction.

## Discussion

The interplay between growth and gene expression is a crucial factor influencing cell population behavior. However, the nature of relationship between this trade-off and the degree of heterogeneity (entropy) within cell populations remains unclear.

Our previous research has identified a link between the growth-gene expression trade-off and cell population heterogeneity, entropy. We found that the switching cost associated with activating burdensome gene circuits is a major driver of cell population dynamics. Specifically, we discovered a burden entropy compensation mechanism that promotes the stability of cell populations upon the activation of burdensome gene circuits. In essence, the loss of growth resulting from burden is compensated by an increase in cellular entropy, leading to the emergence of cells with reduced gene activation levels and, consequently, reduced burden within the population. Our latest results show that the cell population exhibited a robust growth rate, but at the cost of reduced expression of the target gene expression and important cell to cell difference in the level of expression (**Figure 4a**). To address this loss of potential, we explored the possibility of reducing cell population entropy and increasing T7 system activation by increasing the frequency of induction. This control method turned out to be a dead end, we observed mutational escape when increasing the frequency of induction, resulting in the emergence of mutant cells with reduced gene expression (**Figure 4b**). This rapid escape is likely due to the increase in mutation rate associated with the degree of burden experienced by cells^**18**^.

**Figure 4.**
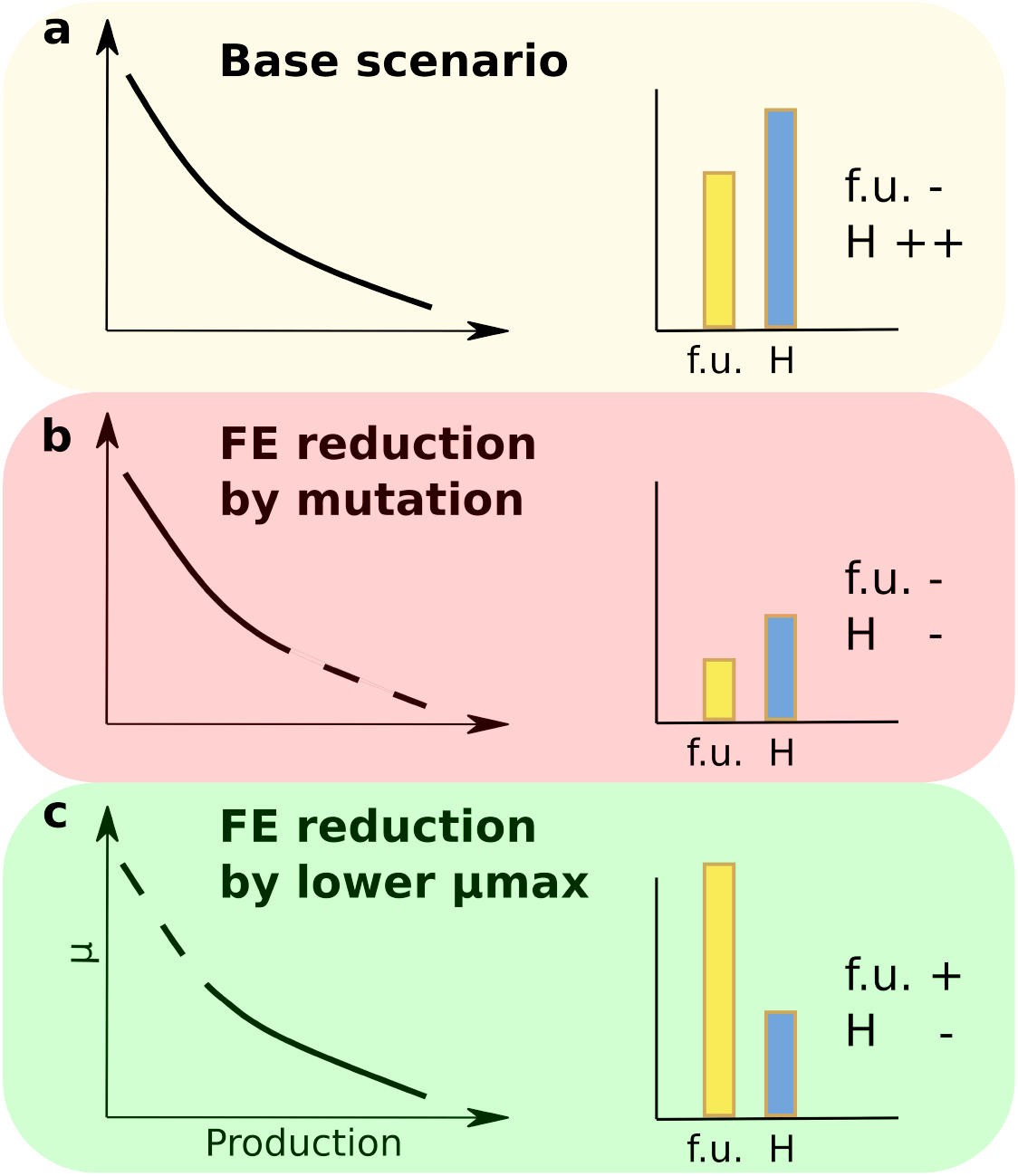
Effect of the growth-production trade-off on the cell population structure: **a** In a traditional continuous cultivation, *E. coli* is grown with glucose, a carbon source providing a high growth rate, and tasked with the production of a burdensome protein. The important switching cost leads to a highly heterogeneous population that harms population productivity. **b** Trying to reduce this heterogeneity by applying periodic pulses can lead to mutational escape, resulting in a population that is more homogeneous but less productive. **c** An alternative way of reducing the switching cost is to reduce the maximum growth rate of the cells, which can be done by lowering the quality of the main carbon source. This enables the creation of a more productive and homogeneous population.

An alternative strategy for mitigating the switching cost is to slow down the maximum growth rate of the population (**Figure 4c**), i.e., limiting the ability of cells with a lower induction to overgrow the producing population. In this study, we employed less effective carbon sources to this end, such as arabinose and xylose. Notably, xylose, which led to the lowest growth rate, was particularly effective in reducing the impact of the switching cost on cell population entropy while increasing the level of T7 system induction throughout the cultivation. Interestingly, the population does not appear to escape control by mutation as fast as it did when we tried controlling it by applying high frequencies of induction. This is likely a result of this softer control method where we did not periodically force harsh inductions but instead reduced that competition between highly induced and less induced cells. Our approach is complementary to the use of “host-aware” gene circuits, which are designed to mitigate in vivo metabolic burden generated by the synthesis and accumulation of various bioproducts^**25**^. Microbial strains engineered to exhibit reduced growth rates could also be explored in the future. Taken together, our approach contradicts the conventional bioprocess design principles, which typically aim to encourage cells to work harder and faster. Our findings reveal that B-E compensation is a critical factor in maintaining the stability of cell populations expressing a burdensome gene. Moreover, a deeper understanding of this phenomenon can lead to the development of innovative strategies for designing more robust continuous bio-process technologies.

## Material and methods

### Strain and growth media

Experiments performed in this work focused on *E. coli* BL21 DE3 with a pET28:GFP plasmid expressing EGFP under the control of P_T7/lacO_ (Plasmid #60733) deposited on Addgene by Matthew Bennett. In all cultivations, the strain was grown in a minimal mineral media containing (in g/l): K_2_HPO_4_ 14.6, NaH_2_PO_4_.2H_2_O 3.6, Na_2_SO_4_ 2, (NH_4_)_2_SO_4_ 2.47, NH_4_Cl 0.5, (NH_4_)2-H-citrate 1, glucose 5, thiamine 0.01, antibiotic 0.1. Thiamine is sterilized by filtration (0.2 mg/l). The medium is supplemented with 3 ml/l of a trace element solution, 3ml/l of a FeCl_3_.6H_2_O (16.7 g/l), 3 ml/l of an EDTA (20.1 g/l) and 2ml/l of a MgSO_4_solution (120 g/l). The trace element solution contains (in g/l): CaCl_2_.H_2_O 0.74, ZnSO_4_.7H_2_O 0.18, MnSO_4_.H_2_O 0.1, CuSO_4_.5H_2_O 0.1, CoSO_4_.7H_2_O 0.21. Filtered sterilized kanamycin (50 mg/l) was added for plasmid maintenance. For bioreactor cultivations, antifoam was added (KS911, 2 drops per liter).

### Bioreactor cultivations and FC analysis

Bioreactor cultivations were performed in a 1 L Bionet F1 lab-scale bioreactor (37°C, 1 VVM, pH 7, and stirring of 1000 revolutions per minute) for the Segregostat cultivation presented in **Figure 1.** All the other cultivations took place in a DASBox (37°C, 1 VVM, pH 7, and stirring of 400 revolutions per minute) with a cultivation volume of 150 ml. The batch was inoculated at an initial optical density (OD) of 0.5 from an overnight preculture (baffled flask, 1 L total volume, 0.1 L working volume). The continuous cultivation was started once oxygen levels began rising at the end of the batch, signaling the transition from exponential to stationary phase.

In the chemostat, the feed contained 5 g/L of the main carbon source (glucose, arabinose, or xylose) and 1 g/L of lactose, the inducer. In the Segregostat, the feed only contained the carbon source (5 g/L glucose) and the inducer was added as a conditional pulse (0.5 g of lactose per pulse). In the experiments where the pulsing frequency varied, each pulse consisted of 1 g of lactose and was automatically triggered from a pre-set .csv file. The automated flow cytometry platform was already described, but briefly: every 12 minutes, a sample is drawn from the bioreactor using a capillary tube and a peristaltic pump. The sample is introduced into an analysis chamber that also contains the flow cytometer. The sample is diluted on the basis of the number of events per microliter (μL) recorded at the previous flow cytometry (FC) analysis by successively flushing and filling the chamber with phosphate-buffered saline (PBS). Then, a FC analysis of 40,000 events is automatically performed, followed by the chamber being cleaned by flushing it multiple times with PBS, and the cycle starts again. In the Segregostat, if more than 50% of the cells are below a fluorescence threshold (1000 arbitrary units on the FL1-A channel), a pulse is triggered.

### Sequencing

At the end of the third cycle of increasingly high pulsing frequency (**Figure 2**), a sample of 1 ml was taken from the reactor and plated to recover 10 random individual colonies. Each colony was then re-grown in LB with kanamycin, the genome extracted and the region of the lac promoter driving the expression of the T7 polymerase was amplified with the following primers:

*FW Sequencing:*

CGCCGTTAACCACCATCAAAC

*RV Sequencing:*

CGCAACTCGTGAAAGGTAG

The same procedure was done from a pre-culture used to start the cultivation to verify that the reactor was not inoculated with a mutant to begin with. All PCR products were then sent for sequencing (Sanger sequencing, Eurofins). All colonies from the pre-culture had the *lacuv5* strong promoter and but 30 % of the one from the end of the cultivations had a lac wt promoter.

*lacuv5*: TATAATGTGTGGAATT

*lacweak*: TAT**GT**TGTGTG**A**AATT

### Maximum growth rate determination

The maximum growth rate under glucose, glycerol, arabinose, xylose, fructose, and mannose was determined by performing batch cultivations of *E. coli* BL21 (DE3) pET28:GFP in mineral media containing 5 g/l of the carbon source and 5 g/l of MOPS buffer to maintain a pH of 7. The batch cultivations were conducted in a flower plate (37°C; 1000 rpm) with a cultivation volume of 1 ml, using a Biolector (Beckman Coulter) instrument, where the back-scatter signal was recorded for 5 wells per condition.

## Software and data availability

All the data presented in this work are available on the following repository: https://zenodo.org/records/13861258. A sample code to treat the data is available on the same repository and utilizes the mBiomass toolbox (https://gitlab.uliege.be/mipi/published-software/mbiomas-core).

## References

(1) Elowitz, M. B.; Levine, A. J.; Siggia, E. D.; Swain, P. S. Stochastic Gene Expression in a Single Cell. Science 2002, 297 (5584), 1183–1186. 10.1126/science.1070919.

(2) Binder, D.; Probst, C.; Grünberger, A.; Hilgers, F.; Loeschcke, A.; Jaeger, K.-E.; Kohlheyer, D.; Drepper, T. Comparative Single-Cell Analysis of Different E. Coli Expression Systems during Microfluidic Cultivation. PLoS ONE 2016, 11 (8), e0160711. 10.1371/journal.pone.0160711.

(3) Ackermann, M. A Functional Perspective on Phenotypic Heterogeneity in Microorganisms. Nat Rev Microbiol 2015, 13 (8), 497–508. 10.1038/nrmicro3491.

(4) Nevozhay, D.; Adams, R. M.; Murphy, K. F.; Josić, K.; Balázsi, G. Negative Autoregulation Linearizes the Dose–Response and Suppresses the Heterogeneity of Gene Expression. Proc. Natl. Acad. Sci. U.S.A. 2009, 106 (13), 5123–5128. 10.1073/pnas.0809901106.

(5) Gardner, T. S.; Cantor, C. R.; Collins, J. J. Construction of a Genetic Toggle Switch in Escherichia Coli. Nature 2000, 403 (6767), 339–342. 10.1038/35002131.

(6) Henrion, L.; Martinez, J. A.; Vandenbroucke, V.; Delvenne, M.; Telek, S.; Zicler, A.; Grünberger, A.; Delvigne, F. Fitness Cost Associated with Cell Phenotypic Switching Drives Population Diversification Dynamics and Controllability. Nat Commun 2023, 14 (1), 6128. 10.1038/s41467-023-41917-z.

(7) Benzinger, D.; Khammash, M. Pulsatile Inputs Achieve Tunable Attenuation of Gene Expression Variability and Graded Multi-Gene Regulation. Nat Commun 2018, 9 (1), 3521. 10.1038/s41467-018-05882-2.

(8) Levine, J. H.; Lin, Y.; Elowitz, M. B. Functional Roles of Pulsing in Genetic Circuits. Science 2013, 342 (6163), 1193–1200. 10.1126/science.1239999.

(9) Sánchez-Romero, M. A.; Casadesús, J. Contribution of Phenotypic Heterogeneity to Adaptive Antibiotic Resistance. Proc. Natl. Acad. Sci. U.S.A. 2014, 111 (1), 355–360. 10.1073/pnas.1316084111.

(10) Arnoldini, M.; Vizcarra, I. A.; Peña-Miller, R.; Stocker, N.; Diard, M.; Vogel, V.; Beardmore, R. E.; Hardt, W.-D.; Ackermann, M. Bistable Expression of Virulence Genes in Salmonella Leads to the Formation of an Antibiotic-Tolerant Subpopulation. PLoS Biol 2014, 12 (8), e1001928. 10.1371/journal.pbio.1001928.

(11) Moreno-Gámez, S.; Kiviet, D. J.; Vulin, C.; Schlegel, S.; Schlegel, K.; van Doorn, G. S.; Ackermann, M. Wide Lag Time Distributions Break a Trade-off between Reproduction and Survival in Bacteria. Proc. Natl. Acad. Sci. U.S.A. 2020, 117 (31), 18729–18736. 10.1073/pnas.2003331117.

(12) Delvigne, F.; Goffin, P. Microbial Heterogeneity Affects Bioprocess Robustness: Dynamic Single-cell Analysis Contributes to Understanding of Microbial Populations. Biotechnology Journal 2014, 9 (1), 61–72. 10.1002/biot.201300119.

(13) David, F.; Berger, A.; Hänsch, R.; Rohde, M.; Franco-Lara, E. Single Cell Analysis Applied to Antibody Fragment Production with Bacillus Megaterium: Development of Advanced Physiology and Bioprocess State Estimation Tools. Microbial Cell Factories 2011, 10 (1), 23. 10.1186/1475-2859-10-23.

(14) Lidstrom, M. E.; Konopka, M. C. The Role of Physiological Heterogeneity in Microbial Population Behavior. Nat Chem Biol 2010, 6 (10), 705–712. 10.1038/nchembio.436.

(15) Xiao, Y.; Bowen, C. H.; Liu, D.; Zhang, F. Exploiting Nongenetic Cell-to-Cell Variation for Enhanced Biosynthesis. Nat Chem Biol 2016, 12 (5), 339–344. 10.1038/nchembio.2046.

(16) Kittler, S.; Kopp, J.; Veelenturf, P. G.; Spadiut, O.; Delvigne, F.; Herwig, C.; Slouka, C. The Lazarus Escherichia Coli Effect: Recovery of Productivity on Glycerol/Lactose Mixed Feed in Continuous Biomanufacturing. Front. Bioeng. Biotechnol. 2020, 8, 993. 10.3389/fbioe.2020.00993.

(17) Kwon, S.-K.; Kim, S. K.; Lee, D.-H.; Kim, J. F. Comparative Genomics and Experimental Evolution of Escherichia Coli BL21(DE3) Strains Reveal the Landscape of Toxicity Escape from Membrane Protein Overproduction. Sci Rep 2015, 5 (1), 16076. 10.1038/srep16076.

(18) Rugbjerg, P.; Myling-Petersen, N.; Porse, A.; Sarup-Lytzen, K.; Sommer, M. O. A. Diverse Genetic Error Modes Constrain Large-Scale Bio-Based Production. Nat Commun 2018, 9 (1), 787. 10.1038/s41467-018-03232-w.

(19) Balázsi, G.; van Oudenaarden, A.; Collins, J. J. Cellular Decision Making and Biological Noise: From Microbes to Mammals. Cell 2011, 144 (6), 910–925. 10.1016/j.cell.2011.01.030.

(20) Eldar, A.; Elowitz, M. B. Functional Roles for Noise in Genetic Circuits. Nature 2010, 467 (7312), 167–173. 10.1038/nature09326.

(21) Nguyen, T. M.; Telek, S.; Zicler, A.; Martinez, J. A.; Zacchetti, B.; Kopp, J.; Slouka, C.; Herwig, C.; Grünberger, A.; Delvigne, F. Reducing Phenotypic Instabilities of a Microbial Population during Continuous Cultivation Based on Cell Switching Dynamics. Biotechnology and Bioengineering 2021, 118 (10), 3847–3859. 10.1002/bit.27860.

(22) Thattai, M.; Van Oudenaarden, A. Stochastic Gene Expression in Fluctuating Environments. Genetics 2004, 167 (1), 523–530. 10.1534/genetics.167.1.523.

(23) Martinez et al. - 2022 - Controlling Microbial Co-Culture Based on Substrat.Pdf.

(24) Kussell, E.; Leibler, S. Phenotypic Diversity, Population Growth, and Information in Fluctuating Environments. Science 2005, 309 (5743), 2075–2078. 10.1126/science.1114383.

(25) Boo, A.; Ellis, T.; Stan, G.-B. Host-Aware Synthetic Biology. Current Opinion in Systems Biology 2019, 14, 66–72. 10.1016/j.coisb.2019.03.001.

(26) Rugbjerg, P.; Sommer, M. O. A. Overcoming Genetic Heterogeneity in Industrial Fermentations. Nat Biotechnol 2019, 37 (8), 869–876. 10.1038/s41587-019-0171-6.

