## Supplementary material for "Lowering the switching cost related to the activation of burdensome gene circuits promotes cell population homogeneity and productivity"

### Supplementary information

#### Supplementary Note 1

To compute the growth rate of the strain as a function of the induction strength (fluorescence content) from FC population snapshots, two assumptions are needed:

1) GFP is not degraded and thus its concentration in the cells only decreases though growth
2) The expression system driving the production of GFP is tight, in other words, during the relaxation phase cells do not produce GFP at all

Both assumptions are reasonable, since: A) GFP is a protein, no degradation tags have been linked to it and the time scale associated to this analysis (only a few hours) is rather small. B) The T7 expression appears really tight as in the batch phase where no lactose is provided, no fluorescence above the auto-fluorescence of a non GFP containing *E. coli* is observed.

From there, the growth rate associated to different level of GFP can be retrieved from multiple FC analysis gathered during the relaxation phase, i.e., when the population fluorescence is decreasing after the increase that followed induction. With these FC files, the fluorescence of the cells can be sorted and characterized with quartiles. These quartiles are collected over a few FC analysis to grasp how their value decreases over time. If the GFP production did not impact the growth, the value of each quartile would drop at a same rate, the population growth rate. Instead of this, what we observe in the case of *E. coli* BL21 (DE3), is that the greater the quartile is, the slower it value decreases over time. A log scale transformation of the quartile values make a simple linear regression sufficient to retrieve the growth rate (the slope of the line) associated with each quartile (Figure 1).

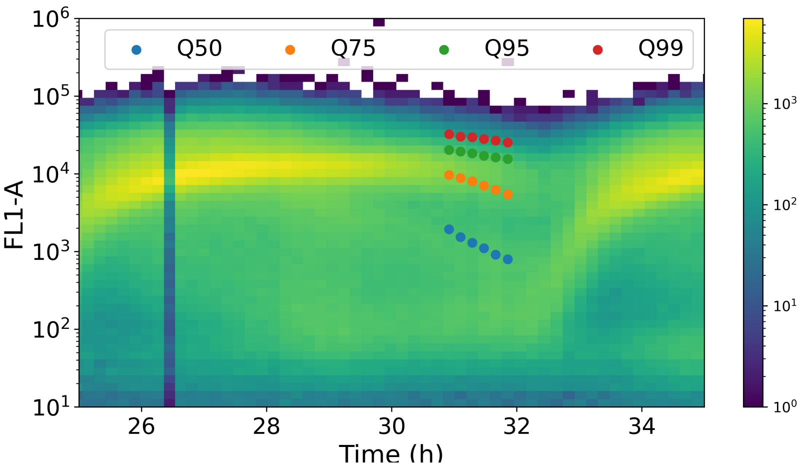


**Figure 1**: Overlapping the quartile values on the density plot during a relaxation phase shows that the slope of the quartile value decrease is inversely proportional to the quartile value.

This method to retrieve the growth rate associated to different GFP content does not work for the low inductions (low quartiles), as the fluorescence is too close to the auto-fluorescence of the cells and thus does not appear to decrease.

#### Supplementary Figure 1

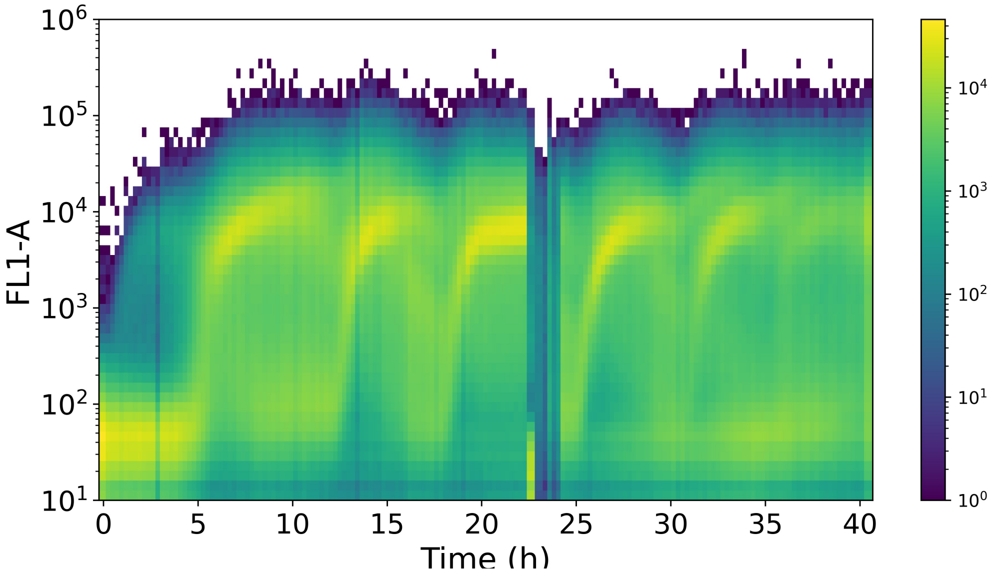


**SI Figure 2:** Time scatter plot of a replicate Segregostat cultivation of *E. coli* BL21 where lactose is added as pulse (0.5 g) once 50% of the population exhibits a fluorescence bellow 1000 fluorescence units (in FL1-A channel)

#### Supplementary Figure 2

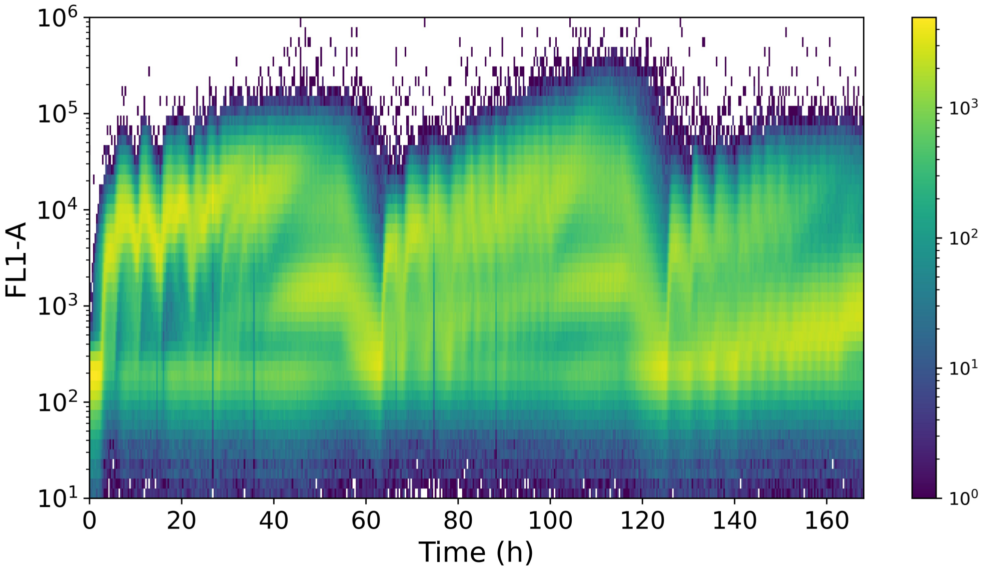


**SI Figure 3:** Time scatter plot of a replicate of the cultivation where lactose is pulsed at an increasingly high frequency three times in a row with 5 retention times in between.

#### Supplementary Figure 3

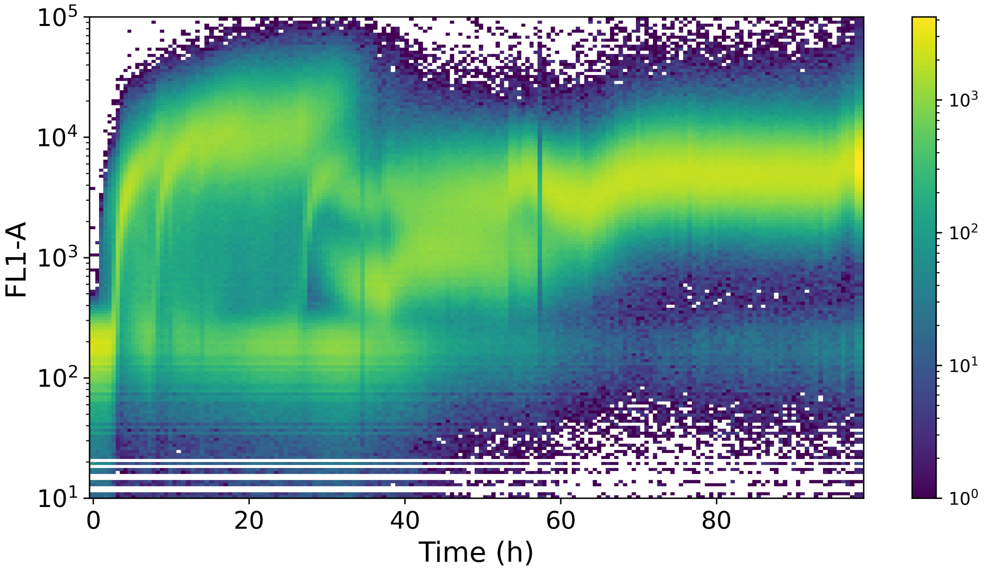


**SI Figure 4:** Time scatter plot of a replicate of a chemostat where the cultivation starts with glucose as the main carbon source, followed with arabinose and then xylose.

#### Supplementary Figure 4

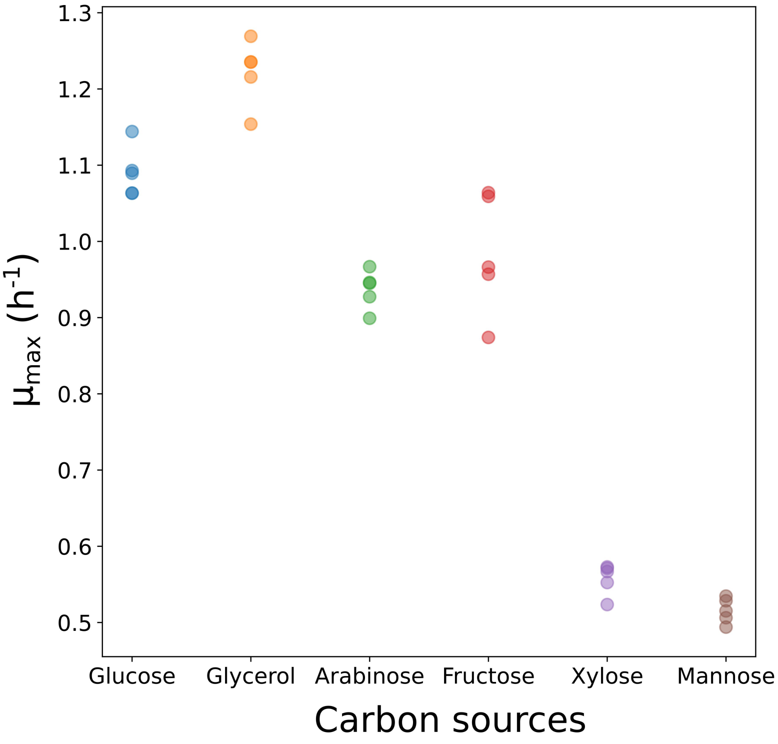


**SI Figure 5:** Maximum growth rate of *E. coli* BL21 (DE3) on multiple carbon sources (n=5). The mean maximum growth rate on glucose, glycerol, arabinose, fructose, xylose and mannose are respectively 1.09, 1.22, 0.93, 0.98, 0.55 and 0.51 $h^{-1}$.
